# PAG-Agent: a biologist-oriented research assistant for context-aware pathway-level analysis and interpretation

**DOI:** 10.64898/2026.06.02.729674

**Authors:** Quang-Huy Nguyen, Zeru Zhang, Duc-Hau Le, Jake Y. Chen, Wei-Shinn Ku, Hao Chen, Zongliang Yue

## Abstract

Pathway analysis is a critical step for translating gene-level omics results into biological mechanisms, yet existing workflows often leave researchers with long lists of statistically significant pathways that are difficult to interpret, validate, and connect to experimental context. We developed PAG-Agent, a biologist-oriented virtual research assistant that integrates pathway-level statistical analysis, context-aware biological interpretation, literature-supported reasoning, and scientific writing support within a unified workflow. PAG-Agent supports bulk and single-cell transcriptomic data and enables users to perform data preprocessing, differential expression analysis, pathway analysis, pathway-level consensus analysis, and pathway-level meta-analysis through click-based and chat-based interactions. Unlike conventional pathway-analysis tools that analyze gene sets largely in isolation, PAG-Agent incorporates experimental conditions and research objectives to prioritize biologically relevant pathways and generate interpretable hypotheses. The system also provides gene and pathway annotation, citation retrieval, visualization, and writing refinement functions. In Alzheimer’s disease case studies using three transcriptomic datasets, PAG-Agent consistently identified neurodegeneration-related pathways across multiple analysis methods and datasets. In citation-retrieval benchmarking, PAG-Agent outperformed six competing LLMs across five common literature-support scenarios, demonstrating improved ability to provide contextually relevant and valid references. Overall, PAG-Agent lowers technical barriers for pathway-level analysis and helps researchers move from transcriptomic data to biologically grounded interpretation, hypothesis generation, and scientific communication.

## 1 Introduction

Advances in high-throughput omics technologies have enabled researchers to profile molecular changes across diverse biological systems and disease conditions. These studies routinely generate lists of differentially expressed genes or gene products from bulk and single-cell experiments. Although informative, gene-level results alone rarely explain the biological mechanisms underlying the observed phenotypes. Pathway analysis (PA) therefore serves as a key step for translating molecular changes into biological processes and functional modules using curated knowledge bases such as KEGG and Gene Ontology (GO) [1–3]. However, PA often produces large collections of statistically significant pathways. Determining which pathways are most relevant to the biological question and integrating them into a coherent interpretation remains a substantial challenge for many researchers.

A wide range of pathway-analysis platforms have been developed to support functional interpretation of both bulk and single-cell omics data. These tools provide capabilities such as differential expression analysis (DEA), PA, consensus analysis, meta-analysis, and visualization [4–17]. Despite these advances, the burden of interpretation remains largely on the user. Researchers must decide which analytical methods to apply, compare outputs across methods or datasets, examine biological relevance, and connect significant pathways to the studied phenotype. These tasks often require substantial domain expertise and manual effort, particularly for early-career researchers and experimental biologists with limited computational backgrounds [18–21]. As a result, obtaining statistically significant pathways does not necessarily translate into efficient biological interpretation.

Large language models (LLMs) have emerged as promising tools for biological interpretation, literature synthesis, and hypothesis generation. Recent studies have demonstrated that LLMs can assist gene-set enrichment analysis by transforming molecular signatures into biologically meaningful functional annotations [22, 23]. However, these approaches typically ignore experimental conditions, research objectives, and other contextual information that motivate the study from the outset. In addition, they provide limited support for the broader workflow that researchers routinely follow to move from omics data to biological insight. Beyond pathway analysis, researchers often need to explore differentially expressed genes, pathways, and biological terms, validate findings through the scientific literature, generate hypotheses, integrate evidence across datasets, and communicate results. Retrieval-augmented generation (RAG) systems have been proposed to improve factual accuracy by incorporating external knowledge during inference [24, 25]. However, these approaches often require substantial computational resources and continuous maintenance of large knowledge repositories. A practical framework that combines pathway-level analysis, context-aware biological interpretation, and reliable literature support within a unified workflow remains lacking.

Here, we present PAG-Agent, a virtual research assistant for pathway-level analysis that integrates statistical analysis, context-aware biological interpretation, literature-grounded reasoning, and research support within a unified workflow. Unlike existing approaches that analyze gene sets in isolation, PAG-Agent incorporates experimental conditions and research objectives throughout the analytical process to prioritize pathways and biological findings that are most relevant to the study context. PAG-Agent supports both bulk and single-cell transcriptomic data and enables researchers to perform data preprocessing, DEA, PA, gene- and pathway-level consensus analysis, and meta-analysis through a user-friendly interface. Beyond statistical analyses, PAG-Agent assists users in exploring genes, pathways, and biological terms, retrieving supporting literature, generating hypotheses, and refining scientific writing. To improve the reliability of literature-supported responses, PAG-Agent incorporates lightweight in-context learning strategies that reduce hallucinations without the computational overhead associated with large retrieval systems. Together, these capabilities enable researchers to move more efficiently from omics data to biological insight.

## 2 Results

### 2.1 PAG-Agent unifies pathway-level analysis and biological interpretation within a single workflow

PAG-Agent simplified the implementation of gene set enrichment analyses by transforming them into intuitive clicks and natural language conversations through its two main modules, including **Click Interaction** (Supplementary Fig. 1) and **Chat Interaction** (Supplementary Fig. 5–29). The Click Interaction module allowed users to upload input data, select preprocessing methods for imputation, normalization, and log2 transformation, choose a differential expression analysis method (limma, edgeR, or DESeq2), and specify a pathway database (KEGG or GO). The Chat Interaction module, powered by GPT-4o-mini (version ‘opt-4o-mini’), served as the core of PAG-Agent. It provided a chat-based interface for users to perform statistical analyses, such as DEA, PA, pathway-level consensus analysis, and pathway-level meta-analysis. Users could also use PAG-Agent to annotate genes or pathways, propose hypotheses linking them to studied phenotypes, refine academic writing, and access scientific citations for validation. Supplementary Fig. 30 presents PAG-Agent’s pipeline as a flow diagram. Based o n t he type o f input data, users can interact with PAG-Agent to perform specific analyses and generate visualizations. Fig. 1 b demonstrates publication-ready graphs generated by the platform. These included gene MA plot, gene volcano plot, pathway volcano plot, pathway bar chart, pathway forest chart, and KEGG pathway maps. The visualizations are available for download in raster (.png) and vector (.svg and .pdf) formats, ensuring compatibility with publications and presentations. Fig. 1c illustrates a simplified dialogue between a human biologist and PAG-Agent for pathway analysis using over-representation analysis (ORA), pathway annotation for Peroxisome (hsa04146), and provision of scientific citations linking Peroxisome and Alzheimer’s disease. Users accessed PAG-Agent through the web portal.

**Figure 1:**
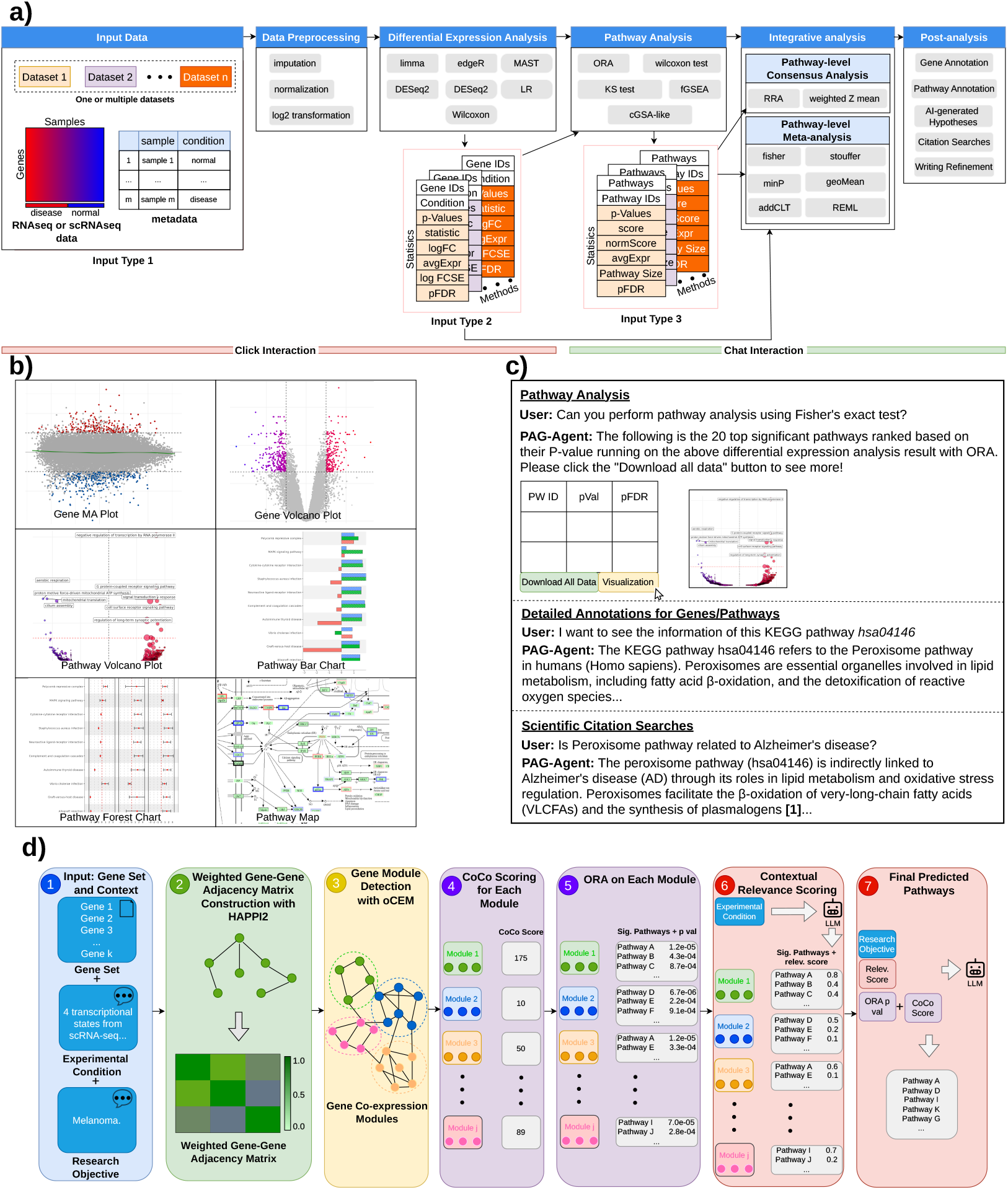
Agentic workflow of PAG-Agent for pathway-level analysis and post-analysis. **a)** PAG-Agent comprises two main modules: **Click Interaction** and **Chat Interaction. Click Interaction**: users can input various input types, including expression matrices with metadata, differential gene expression results, or pathway analysis results. Users can also select data preprocessing methods, choose a differential analysis method (limma, edgeR, or DESeq2), and specify a pathway database (KEGG or GO). **Chat Interaction**: users interact with PAG-Agent through a chat interface to perform four pathway analysis methods, two consensus analysis methods, and six metaanalysis methods. Additionally, users leverage PAG-Agent for post-analysis, including gene or pathway annotation, hypothesis generation linking significant genes or pathways to studied phenotypes, citation searches for validation, and academic writing refinement. **b)** PAG-Agent produces publication-ready graphs as raster (.png) and vector (.svg and .pdf) images. **c)** Simplified example of a dialogue between a human biologist and PAG-Agent during a Chat Interaction session in English (see Supplementary Information Section 2 for details). **d)** Context-aware pathway prioritization using experimental conditions and research objectives to identify biologically relevant pathways from ORA-enriched candidates.

To illustrate PAG-Agent’s capabilities, we analyzed three Alzheimer’s datasets: *GSE153873* [26], *GSE61196* [27], and *GSE5281* [28] (Input Type 1 in Fig. 1a). Detailed descriptions of these datasets were provided in Supplementary Fig. 2 and Supplementary Table 1. We chose the Alzheimer’s datasets because the KEGG pathways for Alzheimer’s disease comprehensively described its mechanisms and biological processes. Furthermore, pathways for Parkinson’s disease, Huntington’s disease, Prion’s disease, and Pathways of neurodegeneration - multiple diseases were known to share genes and mechanisms with Alzheimer’s disease [29–32]. Therefore, we expected these pathways to emerge as statistically significant in our analysis. To further demonstrate PAG-Agent’s ability to perform PA using a single DEA outcome (Input Type 2 in Fig. 1a and Supplementary Fig. 3) and conduct meta-analysis using individual sets of DEA outcomes and PA outcomes (Input Type 2 and Input Type 3 in Fig. 1a, and Supplementary Fig. 4), we generated example DEA and PA result files. Detailed methods for generating these files were provided in the Supplementary Information Section 2.

### 2.2 PAG-Agent supports modality-specific preprocessing and differential expression analysis

Fig. 2 illustrates the user interface of PAG-Agent’s Click Interaction and Chat Interaction modules. In the Click Interaction module, we selected the example dataset *GSE153873-RNAseq*, which meant we used the *GSE153873* datasets consisting of the expression data matrix, metadata, and mapping files. The mapping file served as a reference for gene identifier conversion, allowing PAG-Agent to map source IDs in the input data to a unique list of Entrez Gene IDs. While DEA did not require Entrez Gene IDs, this conversion was a prerequisite for PA and integrative analysis (i.e., consensus analysis and meta-analysis).

**Figure 2:**
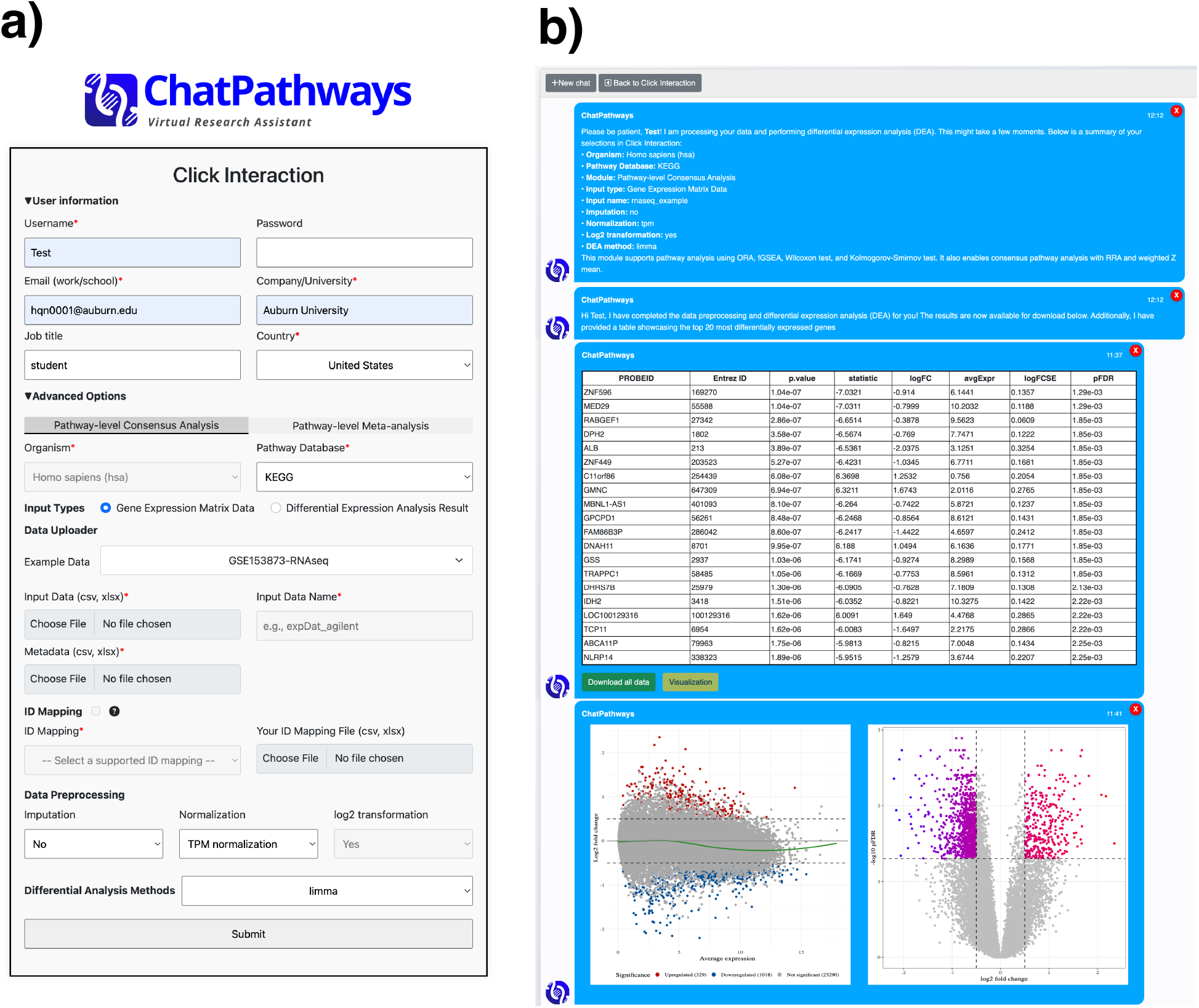
PAG-Agent’s Click Interaction (a) and Chat Interaction (b). **a)** User interface of PAG-Agent’s Click Interaction (see Supplementary Information Section 1 for detailed descriptions of the fields). **b)** After submission, PAG-Agent preprocesses and performs DEA on *GSE153873*. A preview table shows the top 20 differentially expressed genes, ranked in increasing order of FDR-adjusted p-value (pFDR). All outputs are available for download via “Download all data”, providing a ZIP file containing the preprocessed data and DEA result files in CSV format, along with publication-ready visualizations, including MA and Volcano plots, in PNG, SVG, and PDF formats. Visualizations of the analysis outcomes can also be viewed using “Visualization”. An MA plot displays the average expression levels (x-axis) against the log_2_ fold-change (y-axis), with most points centered around zero, indicating no change; deviations suggest differential expression. The colored points are the DE genes with pFDR < 0.05 and absolute log_2_ fold-change > 0.5. A volcano plot shows log_2_ fold-change (x-axis) versus the negative log_10_ p-value (y-axis), highlighting genes with significant expression changes. The colored points are the DE genes with pFDR < 0.05 and absolute log2 fold-change > 0.5.

PAG-Agent provided preprocessing strategies tailored to various expression data types, ensuring compatibility with the specific assay techniques used to generate them (see Methods). For this example, PAG-Agent normalized *GSE153873* (RNAseq) from raw counts to transcripts per million (TPM), applied log_2_ transformation to the normalized data, and performed DEA using limma [33] on the preprocessed dataset (Fig. 2a). For both *GSE61196* (Agilent) and *GSE5281* (Affymetrix), we applied quantile normalization and log_2_ transformation for data preprocessing, followed by limma for DEA. After submitting data, PAG-Agent redirected users to separate chat sessions, each corresponding to a single input. The platform provided a preview table of the DEA results displaying the top 20 differentially expressed genes (Fig. 2b). Further, users could click on “Visualization” to generate the MA and Volcano plots, providing a visual summary of the DEA results. All outputs from this step were available for download via the “Download all data” button, which provided a ZIP file containing the preprocessed data and DEA result files in CSV format, along with the MA and volcano plots in PNG, SVG, and PDF formats. The outcomes of this step for *GSE61196* and *GSE5281* were shown in Supplementary Fig. 6 and 7, respectively.

PAG-Agent visualized a gene network retrieved from KEGG. It mapped DEA results (i.e., log2 fold-change values) onto the network. Users applied this feature to a single dataset or integrate multiple DEA results from different datasets. Supplementary Fig. 30 shows when users could leverage this PAG-Agent’s feature. To illustrate, we used the gene network of the KEGG Alzheimer disease pathway (https://www.genome.jp/pathway/hsa05010). Fig. 3a shows the mapping of log2 fold-change values of differentially expressed genes onto components of the Alzheimer’s disease pathway. Most significant genes belonged to components associated with the PI3K complex, proteasome, and Amyloid *β*. Supplementary Fig. 26 illustrates the integration of the three DEA results from the three Alzheimer’s datasets analyzed using limma into the network.

**Figure 3:**
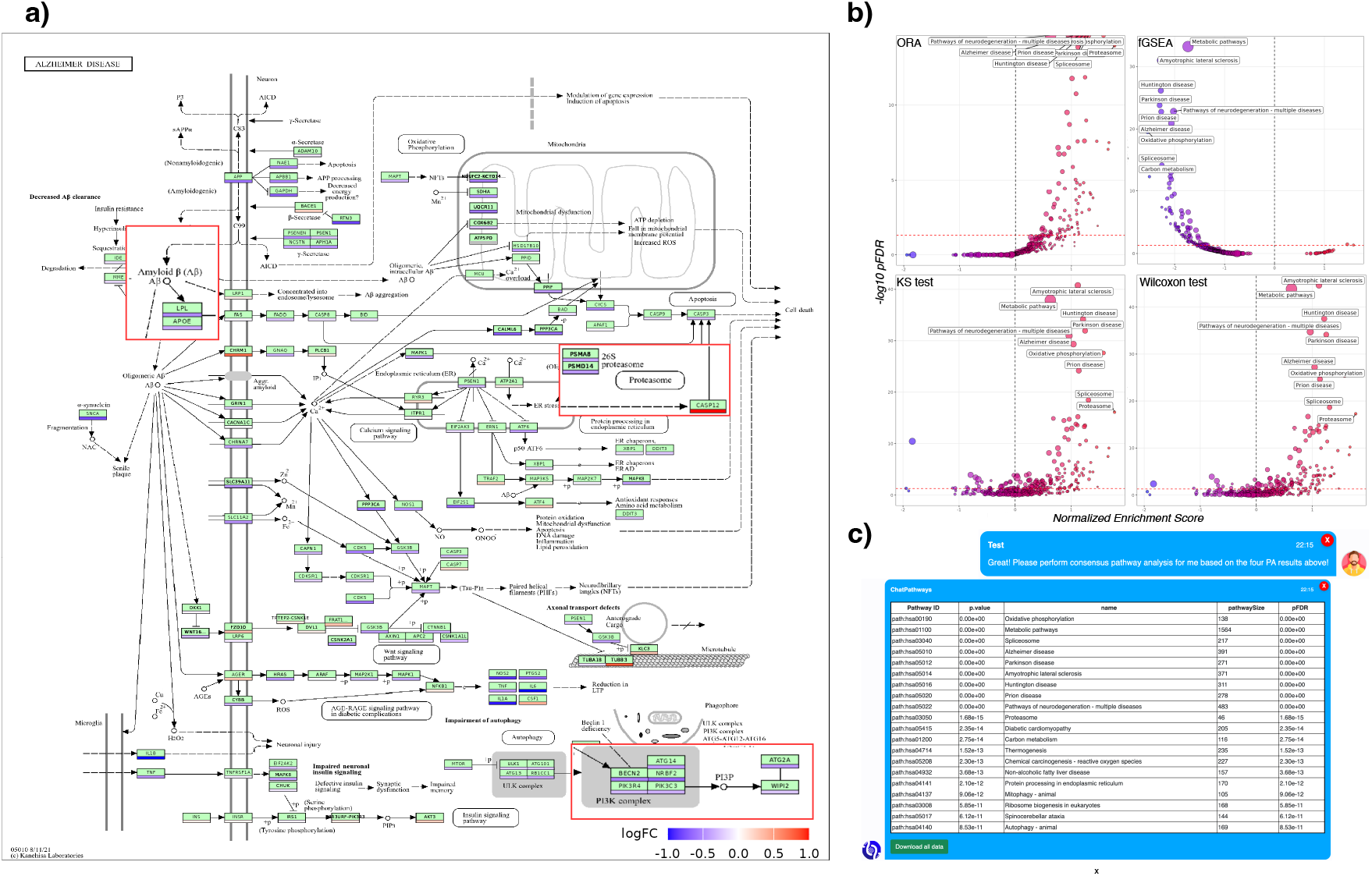
Visualizations of analysis results for *GSE153873*. **a)** Gene network of the KEGG Alzheimer’s disease pathway (hsa05010) with expression changes. Green nodes represent genes involved in the pathway, with bars beneath each node indicating the direction of regulation (up- or down-regulated). **b)** Volcano plots displaying pathway analysis (PA) results for *GSE153873* using over-representation analysis (ORA), fast Gene Set Enrichment Analysis (fGSEA), Kolmogorov-Smirnov (KS) test, and Wilcoxon test. **c)** Pathway-level consensus analysis outcomes using weighted Z mean applied to the four PA results (**b**), presented as a preview table showing the top 20 significant pathways ranked by pFDR. Dialogues between a human biologist and PAG-Agent for reproducing these results are detailed in Supplementary Information Section 2.

### 2.3 PAG-Agent consistently identifies neurodegeneration-related pathways across Alzheimer’s datasets

We continued the current chat session by prompting PAG-Agent to perform PA with KEGG on the three DEA outcomes obtained above, using four available methods: over-representation analysis (ORA), fast Gene Set Enrichment Analysis (fGSEA) [34], Kolmogorov-Smirnov (KS) test, and Wilcoxon test. Each method predicted a list of pathways significantly impacted between the two studied phenotypes (i.e., healthy versus Alzheimer’s disease). As shown in Fig. 3b, pathways associated with neurodegenerative diseases ranked prominently, including Alzheimer disease (hsa05012), Parkinson’s disease (hsa05012), Huntington’s disease (hsa05016), Prion disease (hsa05020), and Pathways of neurodegeneration – multiple diseases (hsa05022). For the *GSE61196* dataset, ORA did not identify any significant pathways (adjusted p-values > 0.05, Benjamini-Hochberg multiple correction [35]), whereas the other methods produced expected results (Supplementary Fig. 12-15). Similarly, for the *GSE5281* dataset, all four PA methods yielded expected results (Supplementary Fig. 16-19).

Next, users were required to perform at least two different PA methods in the chat session before running pathway-level consensus analysis. For *GSE153873* and *GSE5281*, we used all four previously generated PA outcomes. For *GSE61196*, we used three PA outcomes, excluding ORA-based results due to the absence of significant pathways. We selected consensus analysis with weighted Z mean (weightedZMean) [36] as an example (see Methods for a complete list of supported consensus analysis methods). Fig. 3b, Supplementary Fig. 21, and Supplementary Fig. 22 illustrate the top 20 significant pathways with high consensus for *GSE153873, GSE61196*, and *GSE5281*, respectively.

In addition, PAG-Agent supported pathway-level meta-analysis. Users could not perform meta-analysis within the current chat session. Instead, they had to return to the Click Interaction module and select Pathway-level Meta-analysis (Supplementary Fig. 1c). Pathway-level meta-analysis required analysis outcomes as input at the gene or pathway level. For gene-level outcomes, users provided a list of DEA outcomes. For pathway-level outcomes, they provided a list of PA outcomes. To demonstrate this functionality, we asked PAG-Agent to perform meta-analysis using Stouffer’s method [37] (see Methods for a complete list of supported meta-analysis methods). We inputted a set of three fGSEA-based PA outcomes generated earlier for *GSE153873* (RNAseq), *GSE61196* (Agilent), and *GSE5281* (Affymetrix) into PAG-Agent to execute meta-analysis at the pathway level. PAG-Agent simplified the result presentation via Volcano plots (Fig. 4a), Forest plots (Fig. 4b), and Bar charts (Fig. 4c). The complete meta-analysis results for gene- and pathway-level inputs were available in Supplementary Fig. 23 and 24, respectively.

**Figure 4:**
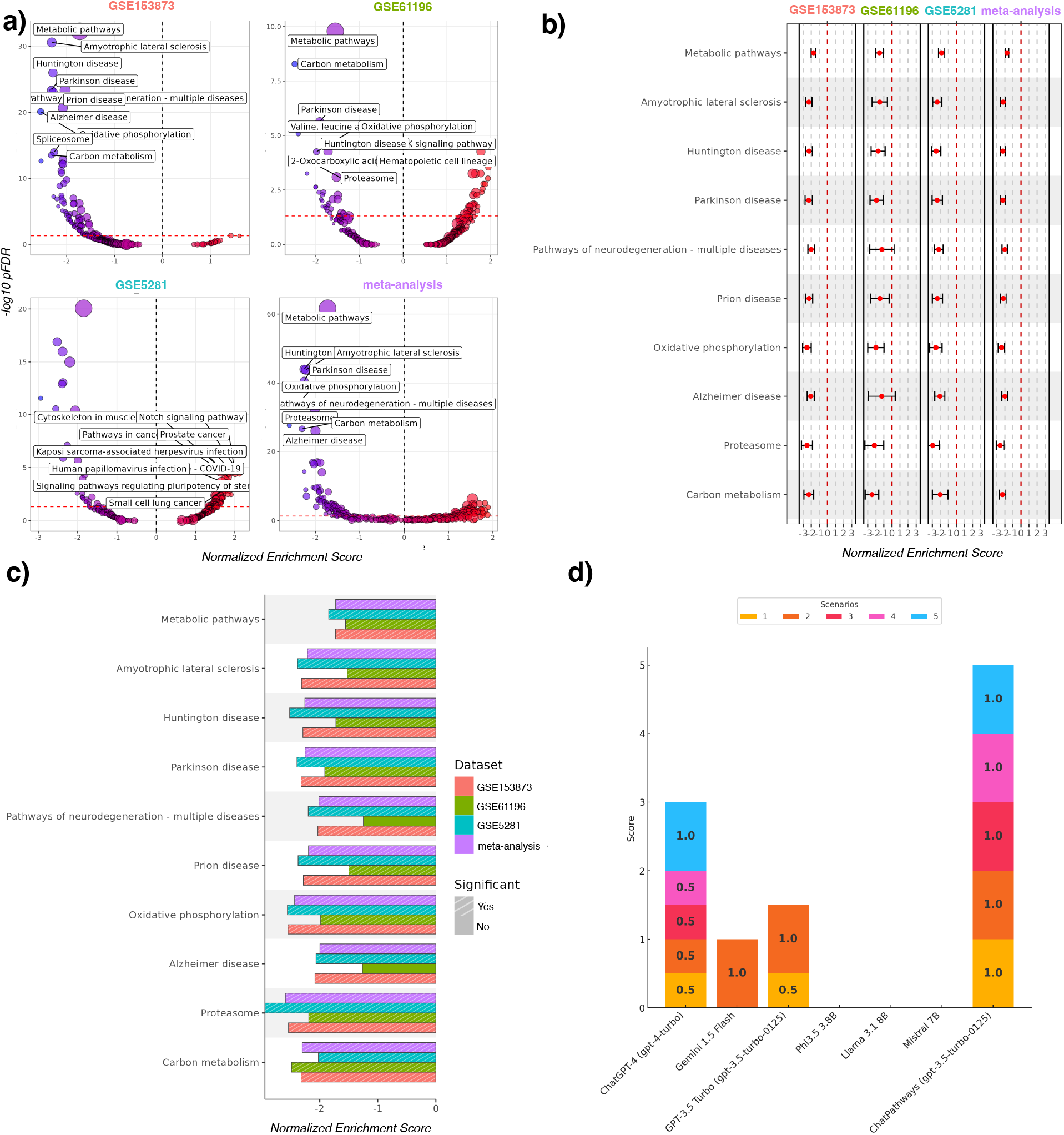
Visualizations of pathway-level meta-analysis results and assessment of PAG-Agent and competing LLMs in providing relevant references. **a-c**) Four Volcano plots, four Forest plots, and bar charts represent the PA results with fast Gene Set Enrichment Analysis (fGSEA) for *GSE153873, GSE61196, GSE5281*, and the meta-analysis result at the pathway level. The dialogues between a human biologist and PAG-Agent to reproduce the results can be found in Supplementary Information Section 2. **d**) Stacked bar chart showcases the capabilities of PAG-Agent and six competing LLMs in providing relevant references across five common scenarios. In the first three scenarios, users seek scientific citations to validate associations between significantly perturbed pathways and specific conditions, with requests expressed in various styles. The fourth scenario tests the ability of the LLMs to retrieve desired information alongside relevant citations. The final scenario challenges the LLMs to incorporate relevant citations into a long, user-written paragraph. (see Methods and Supplementary Table 2 for more details).

In summarize, PAG-Agent performed a total of 16 pathway-related analyses across the three Alzheimer’s datasets. Six pathways associated with neurodegenerative disorders, including Pathways of neurodegeneration - multiple diseases, Alzheimer’s disease, Huntington’s disease, Parkinson’s disease, Amyotrophic lateral sclerosis, and Prion disease, were consistently significant in 15 out of 16 analyses.

### 2.4 PAG-Agent facilitates literature-grounded interpretation and hypothesis generation

Detailed annotations for genes or pathways provided critical biological context for interpreting results and selecting candidates for follow-up studies. Traditionally, computational scientists performing DEA and PA would deliver lists of significant genes or pathways to biologists without additional context. Consequently, experimentalists had to identify relevant knowledge bases, extract, and analyze the required information to address their queries. Common questions included understanding the biological role of a gene or pathway, identifying related pathways, determining which drugs targeted a specific gene or pathway, and exploring associations between genes or pathways and the studied phenotypes. This process was particularly challenging for novice biologists unfamiliar with bioinformatics. For example, in Fig. 2b, *ALB* (Entrez ID 213) ranked high on the DEA results, while PAG-Agent identified metabolic pathways (hsa01100) as significant (Fig. 3). However, even domain experts struggled to immediately provide accurate biological context or justify associations with Alzheimer’s disease. PAG-Agent, leveraging the combined power of GPT-4o-mini and in-context learning techniques (Methods), rapidly generated detailed and accurate annotations of these genes and pathways as well as made hypothesis on their relation with the studied conditions, as shown in Supplementary Fig. 27 and 28. These AI-generated hypotheses provided biologists with interpretable results or actionable starting points for further investigation. Notably, we encouraged LLMs to generate creative and novel hypotheses, exploring plausible connections between discoveries and phenotypes. We tolerated outputs containing hallucinations when they introduced potential insights that had not been previously considered.

A key aspect of comparative studies involved determining whether any previous articles reported similar findings. Supplementary Fig. 29 illustrates PAG-Agent, integrated with a search engine (see Methods), retrieved relevant references linking Oxidative Phosphorylation to Alzheimer’s disease. It further summarized these references and presented them in an academic format suitable for immediate use. To benchmark its performance, we compared PAG-Agent against six competing LLMs across five common scenarios (see Methods and Supplementary Table 2). Importantly, we intentionally downgraded GPT-4o-mini (version ‘gpt-4o-mini’) to GPT-3.5 Turbo (version ‘gpt-3.5-turbo-0125’) as the backend for PAG-Agent to highlight the effectiveness of our proposed strategy for mitigating hallucinations. As shown in Fig. 4d, PAG-Agent consistently outperformed the competing LLMs. Open-weight models, including Phi3.5, Llama 3.1, and Mistral, failed all five scenarios. Scenarios 3, 4, and 5 posed the greatest challenges. While ChatGPT-4 (version ‘gpt-4-turbo’) scored 0.5, 0.5, and 1, respectively, PAG-Agent achieved perfect scores in all three scenarios. Generating a complete response per scenario took PAG-Agent 3–5 seconds, depending on response length and internet speed, comparable to GPT-3.5 Turbo via its API. Notably, GPT-3.5 Turbo without our proposed strategy performed well in only one scenario and moderately in another, achieving a total score of 1.5 out of 5. This result underscored how our strategy substantially reduced AI hallucinations without increasing runtime or computational cost.

English was the global language of science and academic writing [38]. However, non-native English speakers faced significant challenges in manuscript writing, requiring more time and effort [39]. Studies highlighted that these difficulties stem primarily from limited language proficiency [39–41]. Common struggles included difficulties in self-expression, slower writing processes, and restricted vocabulary [40]. These challenges arise not only from linguistic factors such as grammar, vocabulary, and sentence structure but also from meta-linguistic factors, including logical sentence connections, paragraph development, and overall organization. Advancements in internet tools and computational technologies made it easier for researchers to address linguistic issues [42]. However, meta-linguistic challenges remained a significant barrier. Poor English proficiency could also delay research publication [43]. PAG-Agent, powered by advanced LLMs, assisted researchers in refining text and improving overall writing quality. By enhancing clarity and coherence, it promoted equality among researchers, regardless of linguistic background or nationality.

## 3 Discussion

Pathway-level analysis often extends beyond statistical enrichment. Researchers must interpret biological findings, evaluate their relevance to the studied phenotype, validate observations through the literature, integrate evidence across datasets, and communicate results. Existing pathway-analysis platforms provide many of the statistical components required for these tasks, whereas recent LLM-based approaches facilitate biological interpretation. However, these capabilities are typically distributed across separate tools and workflows. PAG-Agent addresses this gap by integrating statistical analysis, context-aware pathway interpretation, literature-supported reasoning, and research assistance within a single platform. The system supports both bulk and single-cell transcriptomic data and provides a unified workflow spanning differential expression analysis, pathway analysis, consensus analysis, meta-analysis, and post-analysis activities. In addition, PAG-Agent incorporates experimental conditions and research objectives into pathway interpretation. The favorable Accuracy, HIT rate, and F1 score achieved across 65 benchmark studies suggest that contextual information can improve the prioritization of biologically relevant pathways beyond conventional enrichment statistics alone.

A central design objective of PAG-Agent is to improve the reliability of literature-supported responses without relying on large, manually maintained knowledge repositories. Retrieval-augmented generation (RAG) systems have emerged as a common strategy for reducing hallucinations by incorporating external knowledge during inference. However, maintaining large collections of scientific materials requires continuous updates and additional computational resources. PAG-Agent instead combines lightweight information retrieval with in-context learning. Relevant references are dynamically retrieved from external sources and incorporated into prompts as contextual evidence for response generation. This strategy enables access to recent scientific findings while maintaining a lightweight architecture. Consistent with this design, PAG-Agent outperformed six competing LLMs in retrieving contextually relevant references across five evaluation scenarios.

Several limitations should be acknowledged. First, our demonstration focused primarily on three Alzheimer’s disease datasets and may not capture the full diversity of biological applications supported by the platform. Second, although the benchmark results support the utility of context-aware pathway prioritization, the benchmark dataset used for evaluation remains relatively modest in size. Future studies will expand benchmarking across additional diseases, organisms, and experimental settings. Future work will also investigate how experimental conditions and research objectives can be incorporated more systematically into pathway interpretation and biological hypothesis generation.

Overall, PAG-Agent lowers the technical barriers associated with pathway-level analysis and interpretation. By combining statistical analyses, biological reasoning, literature support, and scientific communication within a unified workflow, the platform enables researchers to move more efficiently from transcriptomic data to biological insight.

## 4 Methods

### 4.1 Large language model behind PAG-Agent

We integrate the ‘gpt-4o-mini’ version of the OpenAI’s GPT-4o-mini model into PAG-Agent through its well-defined Application Programming Interfaces (APIs). The model allows adjustment of the “temperature” parameter, which controls response variability. Lower temperatures produce more reproducible and conservative outputs [44, 45]. The effect of the temperature parameter on LLM analyses is beyond the scope of this study. We set the temperature to 1.5 for gene and pathway annotation tasks and 0.7 for all other tasks. Annotation tasks involve the biological context of discovered genes and pathways, as well as AI-generated hypotheses on their associations with the studied phenotype. A higher temperature encourages more creative hypotheses from LLMs.

### 4.2 Prompt engineering

The collaborative functioning of eight engineered prompts underpins the smooth operation of PAG-Agent (Supplementary Fig. 31). These prompts included: (i) **User Intent Detection**: This prompt detects user intents, each related to a specific task. (ii) **Pathway Analysis Method Detection**: This prompt identifies the pathway analysis method requested by the user. (iii) **Meta-analysis Method Detection**: This prompt determines the meta-analysis method requested by the user. (iv) **Gene Annotation**: This prompt places a gene to its biological context. (v) **KEGG Pathway Annotation**: This prompt provides information related to a specific KEGG pathway. (vi) **Gene Ontology Term Annotation**: This prompt provides information about a specific Gene Ontology (GO) term. (vii) **Citation Retrieval**: This prompt finds relevant and valid citations based on user-provided context. (viii) **Writing Refinement**: This prompt refines user-submitted academic writing.

We engineer these prompts to identify predefined keywords that indicate user intents for PAG-Agent. **User Intent Detection** includes 10 preset user intents, each linked to a list of keywords. For example, the pathway analysis intent includes keywords: “*Users mention the following keywords: “pathway analysis”, “enrichment analysis”, “functional analysis”, “gene set analysis”, “gene set enrichment analysis”, “fgsea”, “ora”, “over-representation analysis”, “ks”, “ks-test”, “ks test”, “kolmogorov-smirnov”, “kolmogorov-smirnov test”, “wilcox”, “wilcox-test”, “wilcox test”, “wilcoxon”, “wilcoxon-test”, “wilcoxon test”, “hypergeometric test”, “hypergeometric”, “fisher”, “fisher’s exact test”, “fisher’s exact”, or “fisher exact test”*“. When a user submits a query like “*Can you perform <u>ora</u> for me?* “, PAG-Agent detects the keyword “ora” and maps it to the PA intent. The system then proceeds to the **Pathway Analysis Method Detection** prompt to determine the specific method (over-representation analysis in this example). Supplementary Fig. 31 outlines the hierarchical organization of these prompts.

To minimize hallucinations in responses from the **Gene Annotation, KEGG Pathway Annotation, Gene Ontology Term Annotation**, and **Citation Retrieval** prompts, we integrate recent and relevant information through in-context learning strategies into the system prompts. For **Gene Annotation**, we use species-specific annotation packages from Bioconductor (https://bioconductor.org/packages/3.20/data/annotation/; release 3.20.0) to retrieve biological data. For instance, when querying the human gene *DPH2* (Entrez ID: 1802) with a prompt like “*I want to get the information of the gene 1802 within the context of Alzheimer’s disease*”, the R annotation package org.Hs.eg.db fetches data from NCBI (https://www.ncbi.nlm.nih.gov/gene/1802) and provides relevant context for PAG-Agent to generate accurate responses. A similar approach is applied to **KEGG Pathway Annotation** and **Gene Ontology Term Annotation**, using the R packages KEGGREST (version 1.46.0) and GO.db, respectively. For **Citation Retrieval**, we leverage the DuckDuckGo Search API (https://api.duckduckgo.com/), accessed via the Python library duckduckgo search (version 7.1.0). This enables the retrieval of valid citations, reducing hallucinations and enhancing the reliability of responses.

### 4.3 Assessment of PAG-Agent’s capability to provide relevant references

Providing scientific references is crucial for enhancing the reliability of analysis results generated by statistical methods. While LLMs can provide citations, their responses often include hallucinated references. To address this, we design PAG-Agent to rapidly deliver scientific citations that are contextually relevant and valid. To evaluate our strategy for efficient citation retrieval, we intentionally downgrade ‘gpt-4o-mini’ to ‘gpt-3.5-turbo-0125’. Six LLMs are selected for evaluation: GPT-3.5 Turbo (version ‘gpt-3.5-turbo-0125’), ChatGPT-4 (version ‘gpt-4-turbo’), Google’s Gemini 1.5 Flash, Meta’s Llama 3.1 8B, Mistral 7B from the Mistral AI team, and Microsoft’s Phi-3.5 3.8B. We access ChatGPT-4 and Gemini 1.5 Flash through their official platforms (https://chatgpt.com/ and https://gemini.google.com/app), while Llama 3.1, Mistral, and Phi-3.5 are queried via the API endpoint of Ollama (https://ollama.com/). The ‘gpt-3.5-turbo-0125’ version of GPT-3.5 Turbo is utilized through its API. The evaluation involves five common scenarios, including but not limited to Alzheimer’s disease. In the first three scenarios, users seek scientific citations to validate associations between significantly perturbed pathways and specific conditions, with requests expressed in various styles. The fourth scenario tests the ability of the LLMs to retrieve desired information alongside relevant citations. The final scenario challenges the LLMs to incorporate relevant citations into a long, user-written paragraph. The prompt strategy for each situation is detailed in Supplementary Table 2. The system prompt used for the six competing LLMs is: “*You are a useful assistant. Please provide related citations and include detailed summaries for each*”. In contrast, PAG-Agent utilizes this system prompt in combination with relevant references retrieved from DuckDuckGo. Importantly, the tempurature parameter cannot be controlled for ChatGPT-4 and Gemini 1.5 Flash, as they are accessed via their web interfaces.

LLM-generated responses are classified into three categories: ‘fully match’, if all citations are real, valid, and contextually relevant; ‘partially match’, if the response includes real and contextually relevant references but not suitable as scientific citations (e.g., online news or non-peer-reviewed sources, excluding open-access preprint repositories like arXiv or bioRxiv); and ‘mismatch’, if the LLM hallucinates citations or fails to address the task adequately. To enable comparison, scores of 1, 0.5, and 0 are assigned to responses classified as ‘fully match’, ‘partially match’, and ‘mismatch’, respectively.

### 4.4 Software architecture

The PAG-Agent platform architecture comprises front-end and back-end components. The front-end is built with jQuery and JavaScript, featuring UI components for data uploading and browsing, as well as modules for data fetching and operations. The back-end consists of three core components: (i) the Python FastAPI framework, which manages client requests and application logic; (ii) Nginx (https://www.nginx.com), which serves static files and acts as a reverse proxy; and (iii) R analysis workers, which perform data preprocessing, differential expression analysis, pathway analysis, integrative analysis, visualizations, and context retrieval for annotations. Data is stored in MongoDB (https://www.mongodb.com), while Redis (https://redis.io/) supports high-throughput session management and real-time features. This architecture is designed to handle complex biological data analyses with reliability, scalability, and high performance.

### 4.5 Input and data management

The PAG-Agent platform supports bulk and single-cell transcriptomic datasets through three input types: (i) a gene expression matrix (Supplementary Fig. 2), (ii) a set of DEA outcomes (Supplementary Fig. 3 and 4), and (iii) a set of pathway analysis outcomes (Supplementary Fig. 4). All input data can be provided in either .csv (comma-separated) or Excel .xlsx formats. For expression matrix input, the dataset can include two separate files: one for the expression matrix and another for metadata. The metadata file must contain two main columns: the first lists sample identifiers, and the second specifies their corresponding conditions (e.g., normal or disease). The platform also supports conversions from other gene identifiers to Entrez Gene IDs. Users can either select an available mapping file or import a custom mapping file (Supplementary Fig. 2) for this ID-to-Entrez conversion.

PAG-Agent includes a file manager for uploading, managing, and downloading expression data. Users can upload files directly from their local machines. To free storage space, files uploaded by anonymous users are automatically deleted after 24 hours. Registered users, identified by username, password, and email, can save data permanently and access their chat sessions across multiple devices. Each chat session receives a unique identifier. This identifier allows users to access past interactions with PAG-Agent without re-uploading input data or reselecting parameters in the Click Interaction module, provided the input data remains available. This feature ensures the reproducibility of analysis outcomes.

### 4.6 Data preprocessing

Data preprocessing is a critical step for all statistical analyses. Zhang et al. [46] have demonstrated that pathway analysis is highly influenced by the normalization procedures applied to gene expression measurements. PAG-Agent provides three preprocessing options depending on input data.

Imputation methods, including mean, median, and zero imputation, are available to handle missing values by replacing them with the average, median, or zero value of the corresponding gene across all samples. Users should select the most appropriate method based on the dataset characteristics. Mean or median imputation is generally suitable for Affymetrix and Agilent data (continuous data) to account for random missing values or reduce skewness. Zero imputation, in contrast, is often more suitable for count-based transcriptomic data, including bulk RNA-seq and scRNA-seq data, where zero values may reflect biological absence or technical sparsity.

For bulk transcriptomic data, normalization options include TPM normalization and quantile normalization. TPM normalization is recommended for bulk RNA-seq data, whereas quantile normalization is recommended for microarray data. For single-cell RNA-seq data, PAG-Agent provides quality-control filtering based on the number of detected genes, total UMI counts, and mitochondrial gene fraction. After filtering, the platform applies library-size normalization followed by natural-log transformation to improve comparability across cells.

For bulk transcriptomic data, log2 transformation is applied to stabilize variance and reduce skewness. For single-cell RNA-seq data, PAG-Agent applies natural-log transformation following library-size normalization as part of the standard preprocessing workflow.

### 4.7 Parameter setting for differential expression analysis

Differential expression analysis (DEA) is a fundamental approach for identifying genes that are differentially expressed between distinct conditions or phenotypes. Numerous robust techniques have been developed for differential analysis [47–49]. In this work, PAG-Agent supports multiple differential expression analysis methods tailored to the characteristics of bulk and single-cell transcriptomic data. For bulk transcriptomic datasets, the platform provides limma [33], DESeq2 [48], and edgeR [50]. For scRNA-seq datasets, PAG-Agent supports the Wilcoxon rank-sum test, MAST [51], logistic regression, and DESeq2.

Users can select the most appropriate method based on the assay technology and data representation. Limma is typically applied to continuous data, such as microarray expression or TPM-normalized bulk RNA-seq data, whereas DESeq2 and edgeR are suited for count-based bulk RNA-seq data or scRNA-seq data. For single-cell RNA-seq analyses, the Wilcoxon rank-sum test and logistic regression provide robust non-parametric and discriminative alternatives, while MAST explicitly models zero inflation and cellular detection rates commonly observed in single-cell measurements. The primary output of these methods includes a list of genes with their corresponding fold-changes, p-values, adjusted p-values. Adjusted p-values, calculated using the Benjamini-Hochberg procedure [35], are used to identify significantly differentially expressed genes.

### 4.8 Parameter setting for pathway-level analyses

The platform supports several widely used pathway analysis (PA) methods, including ORA, fGSEA [34] (default), the KS test, and the Wilcoxon test, each designed to detect different data patterns. In this work, the default method refers to cases where the User Intent Detection prompt identifies a user intent for PA, but the Pathway Analysis Method Detection prompt cannot determine the preferred method (e.g., due to typos or unspecified method names). In such cases, PAG-Agent defaults to using fGSEA.

PAG-Agent currently supports 10 organisms: *H. sapiens, M. musculus, R. norvegicus, D. rerio, D. melanogaster, C. elegans, S. cerevisiae, A. thaliana, S. pombe*, and *P. falciparum*. Gene sets for these species from KEGG (Release 112.1) and GO (2024-11-03) are integrated into the platform. Previous studies have highlighted the adverse impact of smaller gene sets on pathway analysis performance [46, 52]. Damian and Gorfine [53] attributed this to larger gene variances in smaller gene sets, which affect the accuracy of test statistics for enrichment. To address this, PAG-Agent excludes gene sets containing fewer than 10 genes.

Existing web-based tools for pathway analysis face several limitations: (i) they cannot combine, compare, or contrast results from multiple methods, (ii) they lack the ability to integrate results across datasets, and (iii) they do not support comprehensive visualization of gene networks and expression changes simultaneously. PAG-Agent addresses these shortcomings through two key strategies: pathway-level consensus analysis and pathway-level meta-analysis.

Pathway-level consensus analysis, also known as multi-method analysis, combines results from different PA methods, such as the KS test and fGSEA, applied to a single gene expression dataset. This approach enhances robustness by identifying pathways consistently enriched across methods. PAG-Agent provides two methods for consensus analysis: weighted Z mean (default) and robust rank aggregation (RRA) [54].

Pathway-level meta-analysis, also known as multi-method and multi-cohort analysis, integrates pathway- or gene-level results across multiple datasets, offering a unified perspective and improving reliability. Pathway-level results are generated from (i) a single dataset using different combinations of DEA and PA methods or (ii) multiple datasets from different experiments analyzed with similar or varied approaches. Similarly, gene-level results combines DEA results within the same context. PAG-Agent supports several established meta-analysis methods: Fisher’s method [55], Stouffer’s method [37] (default), minimum p-value [56], geometric mean, addCLT [57], and restricted (residual) maximum likelihood (REML) [58]. These methods are chosen for their ability to account for heterogeneity across studies, yielding more accurate combined statistics.

Importantly, identification through these integrative analyses does not necessarily imply greater biological significance for a specific gene or pathway. Instead, these methods provide a robust foundation for generating hypotheses that can be explored further in experimental studies.

### 4.9 cGSA-like methodology

#### 4.9.1 Methodology

To improve the biological relevance of pathway interpretation, PAG-Agent incorporates a context-aware pathway prioritization workflow. Starting from a set of differentially expressed genes derived from bulk or single-cell transcriptomic analyses, we first construct a weighted gene–gene interaction network using the HAPPI2 database. Each edge is weighted by the corresponding HAPPI confidence score, resulting in a weighted adjacency matrix that captures both the connectivity and confidence of known functional relationships among genes. Next, we apply the oCEM framework to identify overlapping gene modules from the weighted adjacency matrix. Specifically, oCEM uses independent component analysis to decompose the interaction network into partially overlapping functional modules, allowing genes to participate in multiple biological processes. This property better reflects the pleiotropic nature of biological systems compared with non-overlapping clustering approaches. For each identified module, we perform ORA against pathway databases to obtain candidate enriched pathways. Because multiple enriched pathways may represent related or partially redundant biological processes, PAG-Agent further prioritizes these candidates using contextual information available from the original study. Specifically, PAG-Agent leverages experimental conditions to estimate the contextual relevance of each candidate pathway using GPT-4o-mini with a temperature of zero. The model assigns a relevance score that reflects the consistency between the pathway and the experimental context. Subsequently, PAG-Agent integrates three complementary criteria when selecting a representative pathway for each module: (i) pathway statistical significance derived from ORA, weighted by module quality through the CoCo score; (ii) contextual relevance estimated from the experimental conditions; and (iii) consistency with the stated research objectives. Based on these criteria, the model selects a representative pathway that best explains the biological question under investigation.

#### 4.9.2 Benchmark dataset construction

Evaluating context-aware pathway prioritization requires benchmark datasets in which biologically relevant pathways can be linked to experimentally derived gene sets. Existing pathway-analysis benchmarks primarily focus on statistical enrichment performance and rarely provide pathway annotations that can serve as reference interpretations for evaluating context-aware reasoning. To address this limitation, we constructed a benchmark dataset from RummaGEO (https://rummageo.com/), a large collection of gene sets derived from publicly available transcriptomic studies. For each study, we collected the differentially expressed gene set together with its corresponding pathway annotations reported in the original source. We then applied a series of quality-control criteria to retain only studies with sufficient information for evaluating pathway prioritization (see Supplementary Information Section 5). Briefly, studies were required to provide both gene sets and associated pathway annotations, enabling direct comparison between predicted pathways and study-reported biological findings. After filtering, the final benchmark consisted of 65 independent studies spanning diverse biological conditions and experimental settings (Supplementary Table 3). For each study, the differentially expressed genes served as the input to PAG-Agent, while the pathway annotations reported by the original study were treated as reference pathways for evaluation. This benchmark enabled systematic assessment of PAG-Agent’s ability to prioritize biologically relevant pathways using experimental conditions and research objectives.

## Data availability

The three Alzheimer’s disease datasets, GSE5281, GSE61196, and GSE153873, were downloaded from NCBI GEO [59] and used as example datasets in this study. We additionally constructed a benchmark dataset from RummaGEO (https://rummageo.com/), from which 65 studies satisfying the filtering criteria described in Supplementary Information Section 5 were retained for evaluation (Supplementary Table 3).

## Code availability

The complete prompt engineering details and code for reproducing evaluation task results are available on Github hhttps://github.com/aimed-lab/PAG-Agents under the MIT License. The PAG-Agent website is free, open to all users, and does not require login.

## Authors and Affiliations

**Department of Computer Science and Software Engineering, Auburn University, Auburn, AL 36849, United States**

Quang-Huy Nguyen

**School of Information and Communications Technology, Hanoi University of Science and Technology, Hanoi, Vietnam**

Duc-Hau Le

## Author contributions

Q-.H.N and D-.H.L designed the study. Q-.H.N conceived the idea, drafted the manuscript, conducted prompt engineering, developed the web application system (frontend and backend), deployed the web interface, and implemented PAG-Agent pipeline. Q-.H.N. and D-.H.L evaluated the analysis and conducted the scientific review of the LLM outputs. D-.H.L edited the writing and provided feedback on the web app’s UI to improve clarity. All authors approved the final version of this manuscript.

